# Parental CG and CHG methylation variation is associated with allelic-specific expression in elite hybrid rice

**DOI:** 10.1101/2020.10.19.345892

**Authors:** Xuan Ma, Feng Xin, Qingxiao Jia, Qinglu Zhang, Tong Hu, Baoguo Wu, Lin Shao, Yu Zhao, Qifa Zhang, Dao-Xiu Zhou

## Abstract

Heterosis refers to the superior performance of the hybrid over the inbred parental lines. Besides genetic variation, epigenetic difference between the parental lines has been suggested to be involved in heterosis. However, precise nature and extent of parental epigenome difference and reprograming in hybrids governing heterotic gene expression remain unclear. In this work, we analyzed DNA methylomes and transcriptomes of the widely cultivated and genetically studied elite hybrid rice SY63, the reciprocal hybrid, and the parental varieties ZS97 and MH63, of which the high-quality reference genomic sequences are available. We show that the parental varieties display important variation in genic methylation at CG and CHG (H=A, C, or T) sequences. Compared with the parents the hybrids display dynamic methylation variation during development. However, many parental differentially methylated regions (DMR) at CG and CHG sites are maintained in the hybrid. Only a small fraction of the DMRs display non-additive DNA methylation variation which, however, shows no overall correlation with gene expression variation. By contrast, most of the allelic-specific expression (ASE) genes in the hybrid are associated with DNA methylation and the ASE negatively correlates with allelic-specific methylation (ASM) at CHG but positively at CG sites. The results reveal a specific DNA methylation reprogramming pattern in the hybrid rice and point to a role of parental CG and CHG methylation divergence in allelic specific expression that has been associated with phenotype variation and hybrid vigor in several plant species.

**One sentence summary:** Parental CG and CHG methylation divergence is maintained in hybrid and is related to allelic specific expression associated with phenotype variation and hybrid vigor.

## INTRODUCTION

Heterosis or hybrid vigor refers to the phenomenon where hybrids exhibit superior performance in the traits of interest relative to their parental inbred lines. This phenomenon has been widely exploited in crop breeding to augment agricultural productivity. Heterosis requires genetic variation between parental lines (Schnable and Springer, 2013). However, the relationship between genetic distance and heterosis is not straightforward. For instance, hybrid vigor can occur in progeny derived from crosses of genetically similar ecotypes in Arabidopsis (Groszmann et al., 2011). Accumulating evidence indicates that epigenetic difference is also involved in heterosis (Chen, 2013; Groszmann et al., 2013; Dapp et al., 2015). For instance, small interfering RNAs (siRNA) and/or DNA cytosine methylation levels in heterotic hybrids of Arabidopsis (Groszmann et al., 2011; Shen et al., 2012; Zhang et al., 2016), rice (Chen et al., 2010; He et al., 2010; Chodavarapu et al., 2012), maize (Barber et al., 2012; He et al., 2013; Seifert et al., 2018), and tomato (Shivaprasad et al., 2012) vary from their parental lines. It has been shown that the RNA-directed DNA methylation (RdDM) pathway, which involves siRNAs, is required for differential DNA methylation in Arabidopsis hybrids (Greaves et al., 2012; Greaves et al., 2014; Zhang et al., 2016).

In plants, DNA cytosine methylation occurs in the context of CG, CHG and CHH (where H is A, C, or T). The major role of DNA cytosine methylation is to silence transposable elements (TE) and repetitive sequences and repress gene promoter activity. CG methylation is maintained during cell division by DNA methyltransferase1 (MET1) by recognizing at methylating hemi-methylated CG sites in the newly replicated DNA. Non-CG (i.e. CHG and CHH) methylation in heterochromatin is maintained respectively by plant-specific Chromomethylases 3 (CMT3 for at CHG sites) and CMT2 (at CHH and CHG sites), which is recruited to heterochromatin by interacting with the histone methylation mark H3K9me2, while CHH methylation in euchromatin regions is maintained by the RdDM pathway involving the Domains Rearranged Methyltransferase2 (DRM2) and siRNA (Law and Jacobsen, 2010). DECREASE IN DNA METHYLATION1 (DDM1) is a nucleosome remodeler, the *ddm1* mutation led to >70% reduction in DNA methylation, predominantly affecting methylation at CG and CHG contexts in Arabidopsis and rice (Zemach et al., 2013; Tan et al., 2016). It was shown that hybrids between Arabidopsis C24 and Columbia-0 (Col) defective in RNA polymerase IV (Pol IV, required for siRNA production) or MET1 function did not reduce the level of heterosis of biomass. By contrast, hybrids with *ddm1* mutation displayed a decreased heterosis level (Kawanabe et al., 2016; Zhang et al., 2016), suggesting that specific DNA methylation pattern and/or levels regulated by DDM1 plays a role in heterosis in Arabidopsis.

It has been suggested that epigenetic systems may be involved in the alteration of gene expression in hybrids that, in turn, could contribute to the hybrid phenotype (Greaves et al., 2015). Work in Arabidopsis indicates that locus-specific DNA methylation divergence between the parental lines can directly or indirectly trigger heterosis (Lauss et al., 2018). Some of the changes in DNA methylation in hybrids correlate with changes in transcription levels, but there is no consistent relationship among the changes in DNA methylation, transcription, siRNA, and the generation of the heterotic phenotype (Crisp et al., 2020). It has been shown that gene allelic-specific expression (ASE) can lead to phenotype variation relevant to heterosis (Springer and Stupar, 2007; Paschold et al., 2012; Goff and Zhang, 2013; Shao et al., 2019). Although genetic difference may contribute to allelic-specific expression, epigenetic mechanism has been shown to be essential in differential expression of parental alleles of imprinted genes in endosperm in plants (Gehring, 2013). However, whether parental epigenome differences and their interaction in hybrid are involved in ASE is unclear. In addition, the precise nature and extent of parental epigenetic difference involved in heterosis and whether specific epigenetic variation can be considered as indicator to predict heterosis in agriculturally important crops remain unknown.

Rice is one of the most important food crops in the world. Rice cultivated in Asia can be divided into two subspecies: *Oryza sativa* subsp. *Indica* (*Xian*) and *O. sativa* subsp. *Japonica* (*Geng*). *Indica/Xian* rice can be further genetically subdivided into two major varietal groups, *indica* I and *indica* II, which have been independently bred and widely cultivated in China and Southeast Asia, respectively (Xie et al., 2015). Hybrids between these groups usually generate strong heterosis. For example, the elite hybrid Shanyou 63 (SY63) from the cross between Zhenshan 97 (ZS97, *indica* I) as female and Minghui 63 (MH63, *indica* II) as male exhibits superiority for a large array of agronomic traits and has been the most widely cultivated hybrid in China for approximately three decades (Zhang et al., 2016; Xie and Zhang, 2018). This particular hybrid rice system has been used as a model system for heterosis study, which has led to identification of a number of genetic loci/genes involved in hybrid vigor relevant to yield and yield traits (Yu et al., 1997; Hua et al., 2003; Huang et al., 2010; Zhou et al., 2012; Huang et al., 2015; Xie et al., 2015). The availability of high-quality reference genome sequences of ZS97 and MH63 and the detailed comparative annotation and analysis of the two genomes and transcriptomes (Zhang et al., 2016; Shao et al., 2019) provide unique opportunity to analyze epigenetic basis of hybrid vigor of the elite intra-subspecific hybrid rice.

In this work, we investigated DNA methylation difference between ZS97 and MH63 and parental methylation interaction in the reciprocal hybrids in shoot and panicle. Our data reveal important divergence at CG and CHG methylation pattern and dynamics between the parental lines which is associated with the allele-specific expression in the hybrids.

## RESULTS

### DNA methylation differences between shoot and panicle of two parents and hybrids

To evaluate DNA methylation difference between parental lines of the elite hybrid rice SY63, we analyzed the DNA methylomes of seedling shoot (at 4-leaf stage) and young panicle (2 mm length, inflorescence meristem) of ZS97 and MH63 and the reciprocal hybrids SY63 and MH63/ZS97 (MZ, with MH63 as female and ZS97 as male). Phenotypically the reciprocal hybrids showed similar growth vigor and other traits including grain yield (Supplemental Fig. S1A). The sequencing depth of the BS-seq data was about 18 to 40 × genome coverage and methylation measurements between the two biological replicates for each genotype were highly correlated (Pearson’s *r* was about 0.97 to 0.99) (Supplemental Table S1, Supplemental Fig. S1B). The overall DNA methylation levels in panicle were higher than in shoot of all genotypes (Supplemental Table S1, Supplemental Fig. S1C). The increase of methylation was consistent with the higher expression of DNA methyltransferases CMT3, DRM2 and MET1-2 in panicle (Supplemental Fig. S2A). Density plots (see Methods) indicated that non-CG methylation levels were largely augmented in panicle versus shoot, whereas CG methylation was not changed as much (Fig. 1A). The increased of non-CG methylation was detected in genes and TEs (Fig. 1B and 1C). Differential methylation regions (DMRs) were tested using the criteria of methylation difference at CG >0.7, CHG >0.5, and CHH >0.2 within 200 bp bins that showed at least 25 informative sequenced cytosines in both samples (see Methods). The analysis revealed that at CG context there were very few hyper (0-2) or hypo (42-55) DMRs between shoot and panicle in the 3 genotypes (Fig. 1D). At CHG context, higher numbers of shoot versus panicle DMRs were found in ZS97 (hypo 4630 and hyper 2517) than MH63 (3 hyper and 496 hypo) and SY63 (68 hyper and 1444 hypo) (Fig. 1D). At CHH sites 42,000-55,000 hypo but only 274-588 hyper DMRs were detected in the 3 genotypes (Fig. 1D). The analysis indicated a predominant increase of CHH methylation during shoot to panicle development and a more dynamic CHG methylation in ZS97 than MH63. The data corroborated previous observation that methylation increases during development in rice (Zemach et al., 2010; Higo et al., 2020), and revealed important difference in CHG methylation dynamics between the two parental lines.

**Figure 1.**
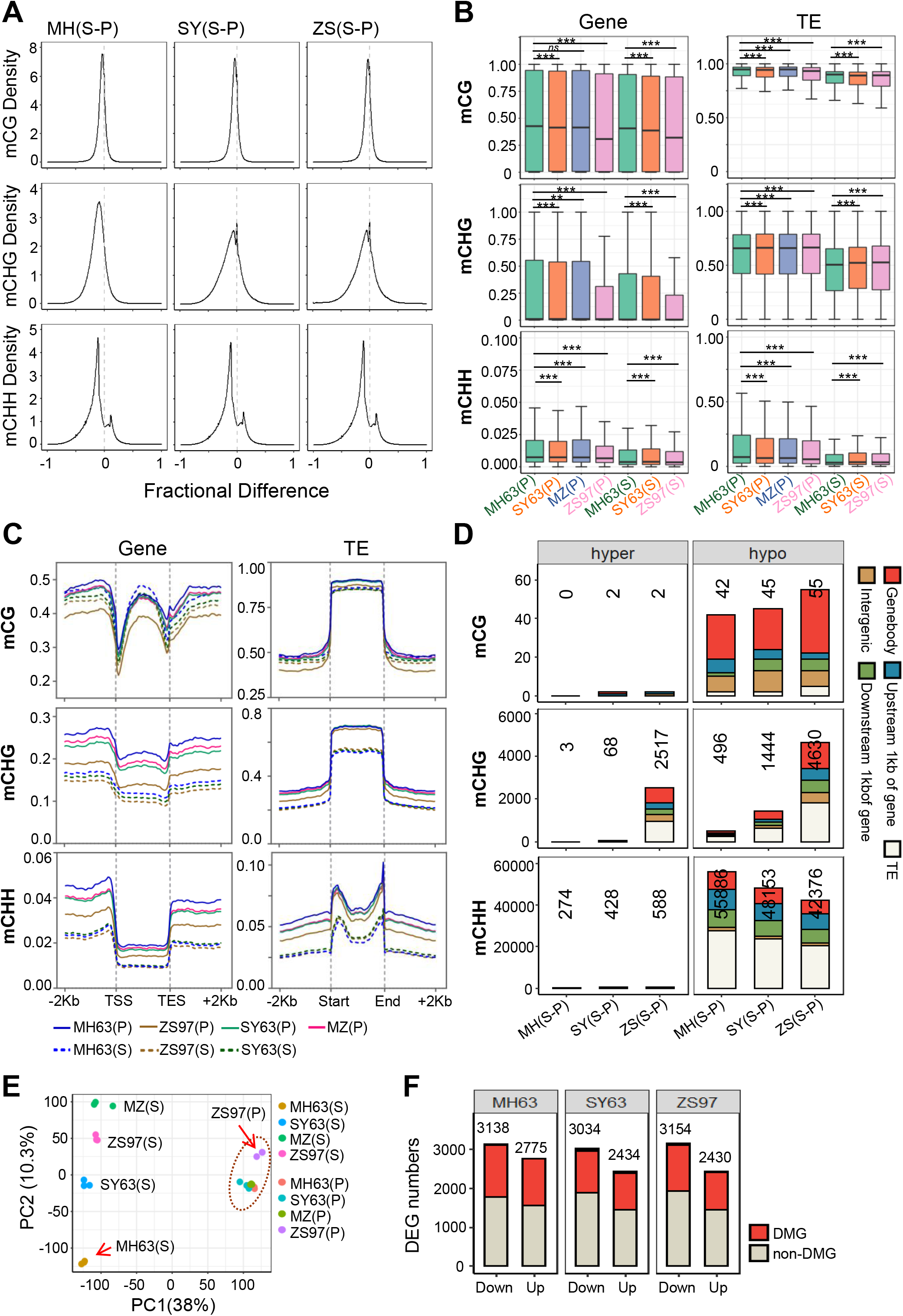
DNA methylation levels in shoot and panicle of MH63, ZS97 and the hybrids SY63 and MZ. (A) Density plots showing the frequency distribution of CG, CHG, and CHH methylation difference at 200 bp windows between shoot and panicle of MH63, ZS97 and SY63. (B) Boxplots showing overall DNA methylation levels (at CG, CHG, and CHH context) in gene and TE of MH63, ZS97, SY63 and MZ in shoot (S) and panicle (P) (** P < 0.01,*** P < 0.001, *ns*, not significant, Wilcoxon rank-sum test). (C) Metaplots showing the average methylation levels within each 50 bp interval plotted in gene and TE of MH63, ZS97, SY63 and MZ in shoot (S) and panicle (P). (D) Numbers of differentially methylated regions (DMRs) of 200 bp bins between shoot and panicle in MH63, ZS97 and SY63, respectively. Different colors showing the DMRs distributed in gene body, upstream 1kb of gene, downstream 1kb of gene, TE and intergenic regions. (E) Principal Component Analysis (PCA) of transcriptomes of MH63, ZS97, SY63 and MZ in shoot (S) and panicle (P). (F) Differentially expressed genes (DEGs) (Q value < 0.01, |log2(Fold Change)| >2) in shoot versus panicle of MH63, ZS97 and SY63. Numbers of DEGs with or without DMRs in gene body or gene flanking regions are indicated.

In parallel, we analyzed the shoot and panicle transcriptomes of the three genotypes. Three biological replicates were sequenced and there were high correlations of the transcript numbers between the replicates (*r* = 0.88 to 1.00) (Supplemental Table S2; Supplemental Fig. S3; Fig. 1E). Principal component analysis indicated that the shoot transcriptomes were distally apart from those of panicle of the different genotypes. The shoot transcriptomes of the hybrids were closer to that of ZS97 while the panicle transcriptomes of the hybrid were closer to that of MH63, which is consistent with the ZS97-like growth vigor at seedling stage and MH63-likely panicle architecture at mature stage observed in SY63 (Xie and Zhang, 2018). In total, 2430 to 3154 up and down-regulated (fold change > 4 ×, Q value < 0.01) genes were detected in shoot versus panicle. About 40% of the DEGs were associated with DMRs (Fig. 1F), suggesting that differential DNA methylation is involved in differential expression in shoot and panicle.

### DNA methylation difference among the parental lines and hybrids

Density plots analysis revealed variation of DNA methylation levels at all sequence contexts between the two parent lines (Fig. 2A). In shoot, the CHH methylation variation was relatively low compared with CG and CHG methylation but became more important in panicle (Fig. 2A). Analysis of DMR between ZS97 and MH63 revealed 4084-6182 hyper and 6959-8028 hypo CG DMRs and 3357-3592 hyper and 3043-3306 hypo CHG DMRs in shoot and panicle (Fig. 2B). More than 72% of the DMR at CG sites and 55% at CHG sites in panicle overlapped with those detected in shoot (Fig. 2C and 2D). The data indicated that the parental lines had substantial difference in CG and CHG methylation, most of which persisted during shoot to panicle development (Fig. 2C and 2D; Supplemental Fig. S4). By contrast, difference at CHH methylation between ZS97 and MH63 was relatively low in shoot (646 hyper and 466 hypo DMRs) and increased in panicle (1650 hyper and 6104 hypo DMRs in ZS97 versus MH63) (Fig. 2A and 2B). The higher number of hypo than hyper DMRs in ZS97 panicle was consistent with the larger increase of panicle CHH methylation in MH63 (Fig. 1D). Nevertheless, methylation variation at CHH sites (especially in shoot) was unexpectedly low between the parental lines. By comparison, the reciprocal hybrids showed 1796 hypo and 1943 hyper CHH DMRs in SY63 compared to MZ with much less or no methylation difference at CG and CHG contexts (Fig. 2A and 2B). The very low divergence of CG and CHG methylation between the reciprocal hybrids suggests that there is little effect of the parent-of-origin in genome-wide methylation pattern at the sequence contexts.

**Figure 2.**
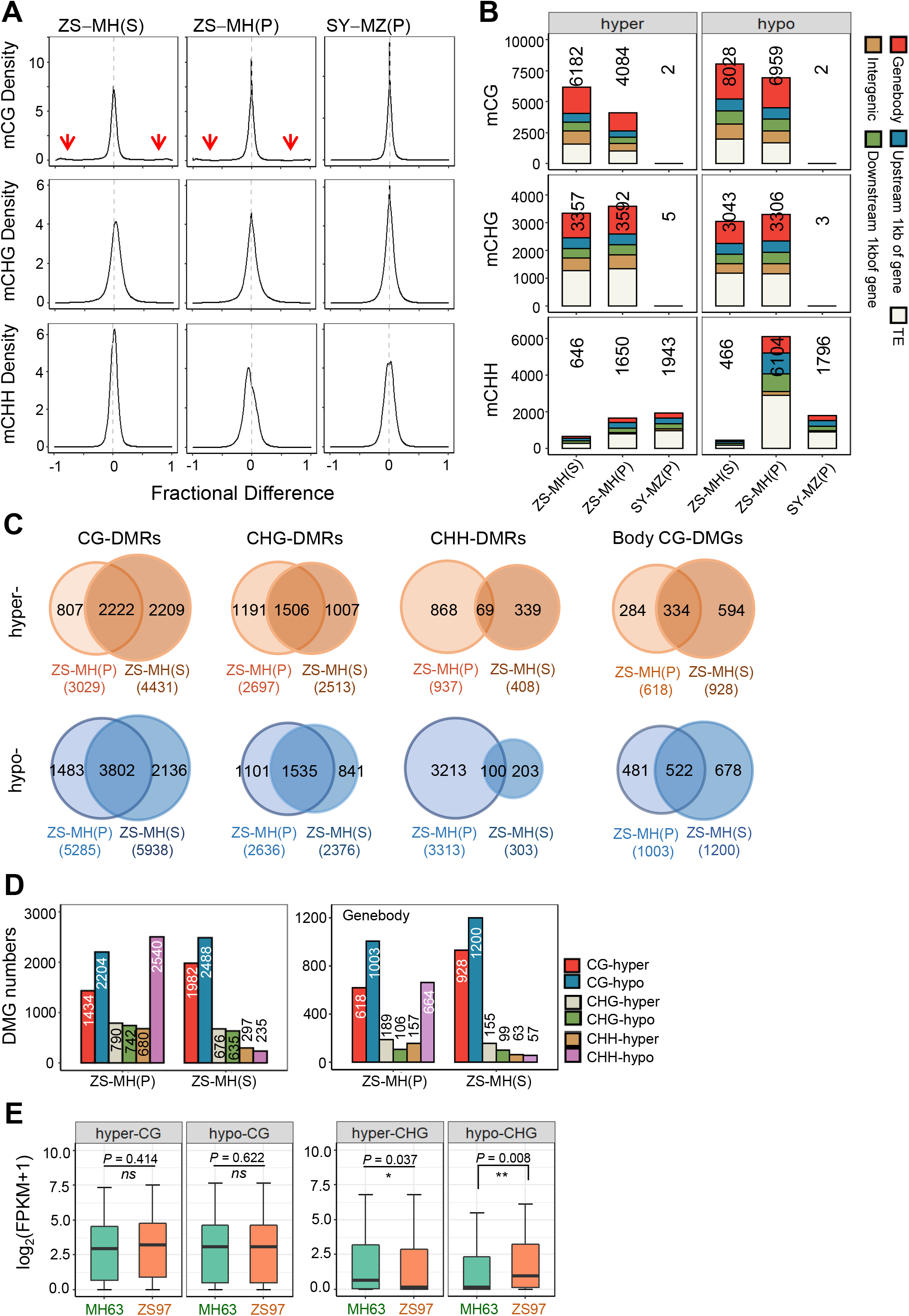
Divergence of DNA methylation landscape between MH63 and ZS97. (A) Density plots showing the frequency distribution of methylation difference at 200 bp windows between MH63 and ZS97 in shoot and panicle, and methylation difference between SY63 and MZ in panicle, respectively. Red arrows indicate small numbers of bins with large difference in CG methylation between ZS97 and MH63. (B) Numbers of ZS97 versus MH63 shoot (S) and panicle (P) DMRs of 200 bp bins and SY63 versus MZ panicle (P) DMRs. Different colors showing the DMRs distributed in gene body, 1kb flanking regions of gene, TE and intergenic regions. (C) Venn diagram showing overlapping DMRs (in CG, CHG, and CHH context) between shoot and panicle. (D) Numbers of differentially methylated genes (DMGs) between ZS97 and MH63 in shoot (S) and panicle (P). Left, genes with the flanking regions; right, gene body only). (E) Boxplots showing the expression level of hyper- or hypo-DMGs (ZS97-MH63) with methylation sites in gene body regions in MH63 and ZS97 panicle. *P* < 0.05 indicates statistical significance (two-tailed *t* test).

Albeit that CG and CHG methylation mainly target TEs and repetitive sequences in the rice genome (Tan et al., 2016), more than 60% of the CG and CHG DMRs between ZS97 and MH63 were detected in genes (body and flanking regions) (Fig. 2B and 2D; Supplemental Dataset S1 and S2). In panicle, there were 1434 and 2204 genes with respectively hyper and hypo CG methylation and 790 and 742 genes with respectively hyper and hypo CHG methylation detected in ZS97 versus MH63 (Fig. 2D). The data highlighted that important variations of genic CG and CHG methylations exist between ZS97 and MH63, which is maintained in most cases during shoot to panicle development.

Many CG and, to a lesser extent, CHG DMRs were located within gene body (Fig. 2B). In fact, nearly 45% of the CG DMGs were gene body CG DMGs (Fig. 2C and 2D; Supplemental Fig. S4). It was shown that gene body CG methylations were associated with constitutively expressed genes with moderate to high expression levels in plants (Bewick et al., 2016). To study whether differential gene body CG methylation is correlated with differential gene expression, we plotted transcript levels of the DMGs between ZS97 and MH63. The analysis revealed no significant difference of expression (Fig. 2E). By contrast, gene body CHG DMGs display a reverse correlation with expression levels between ZS97 and MH63 (Fig. 2E).

There are 1,284,423 single nucleotide polymorphisms (SNPs, not including InDels) between MH63 and ZS97 genomes (Zhang et al., 2016). Both SNP and non-SNP bins showed methylation difference between ZS97 and MH63 (Supplemental Fig. S5A), but percentages of DMR in SNP bins are higher than those in non-SNP bins (Supplemental Fig. S5B).

### Differential methylation in the hybrids relative to the parents

Overall methylation levels in the reciprocal hybrids SY63 and MZ were intermediate between the parents (Fig. 1B; Supplemental Fig. S1C; Supplemental Table S1). Density plots revealed that in shoot, CG and CHG methylations show clear variations between SY63 and the parents but the difference became less pronounced in panicle (Fig. 3A). Conversely, the variation of CHH methylation in shoot was lower than that of CG or CHG methylations, but became more pronounced in panicle (Fig. 3A). This likely reflects the dynamic change of the methylation divergences from shoot to panicle between the parents (Fig. 2A and 2B). Similar profiles of density plots are observed in panicle methylation of the reciprocal hybrid MZ relative to the parents (Fig. 3A), consistent with the low methylation divergence between SY63 and MZ (Fig. 2A and 2B). The analysis revealed substantial DNA methylation differences in the hybrids compared to either of the parents, which became more or less pronounced in panicle depending on the sequence context.

**Figure 3.**
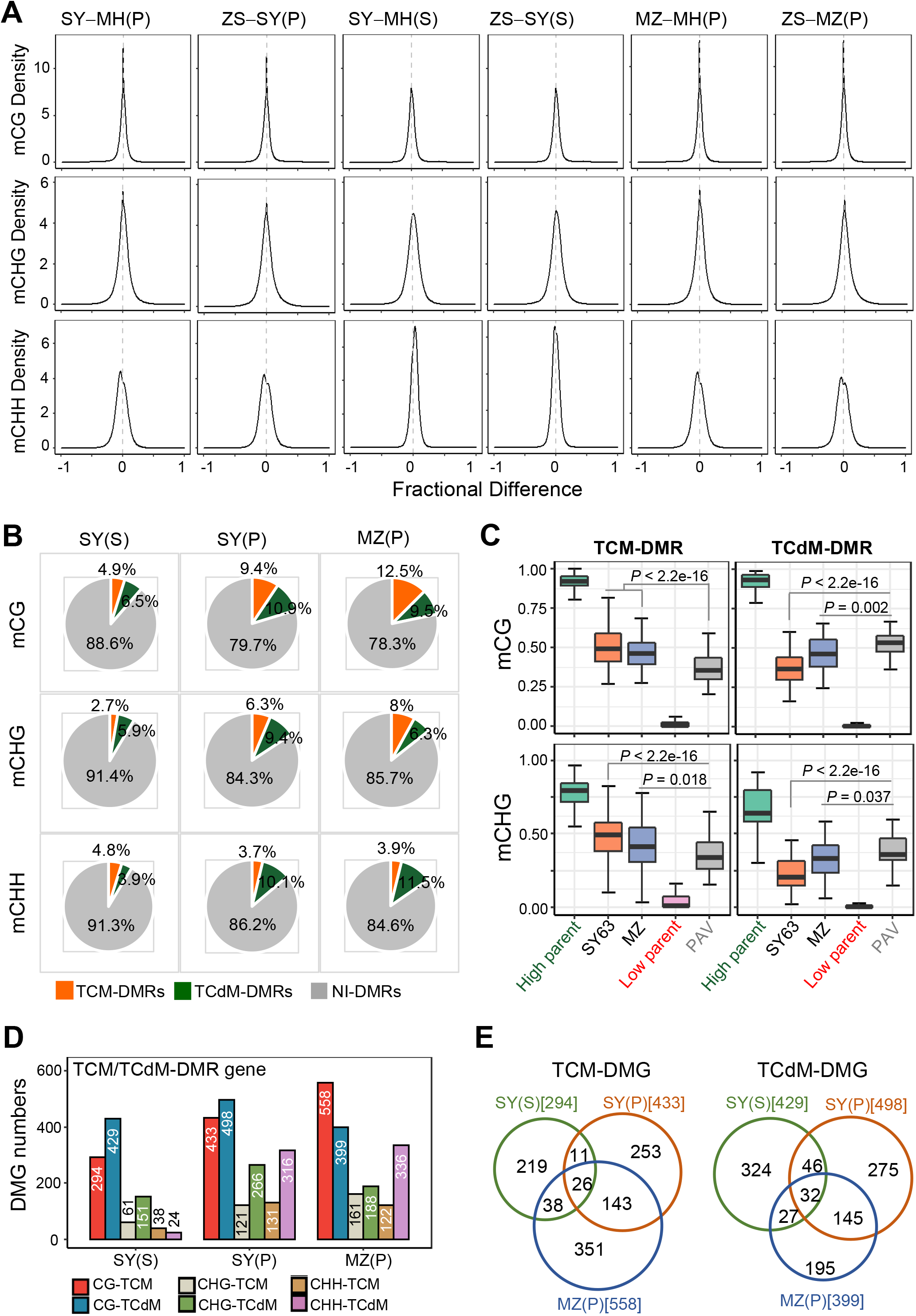
Methylation variation and interaction of parental DMRs in the hybrids. (A) Density plots showing the distribution of methylation difference frequency at 200-bp windows between the hybrids SY63 and MZ and either of the parents in shoot and/or panicle. (B) Percentage of the TCM-DMR, TCdM-DMR and NI-DMR regions in SY63 shoot and panicle and MZ panicle. (C) Boxplots showing CG and CHG methylation levels of TCM-DMR, TCdM-DMR in SY63 panicle, MZ panicle and the parents. (D) Numbers of TCM-DMG (differentially methylated genes), TCdM-DMG and NI-DMG in SY63 shoot, SY63 panicle and MZ panicle. (E) Venn diagram of overlapping TCM-DMGs or TCdM-DMGs in CG context between SY63 shoot, SY63 panicle and MZ panicle.

To identify differential methylation between hybrids and the parents, we calculated the predicted additive value (PAV) of the DMRs identified between the parents as previously described (Schultz et al., 2012; Zhang et al., 2016) (see Methods), and compared the methylation levels of hybrid with the PAVs in corresponding regions (within 200 bp bins). The hybrid bins (F1) that show >PAV and FDR <0.01 are known as trans-chromosomal methylation regions (TCM), whereas bins with F1 < PAV and FDR <0.01 are considered as trans-chromosomal demethylation regions (TCdM) (Zhang et al., 2016) (Supplemental Fig. S6A). These differential methylations are supposed to be from non-additive interaction, while the remaining bins are resulted from additive interaction between the parental methylation (Zhang et al., 2016). In shoot, only a small portion of the parental DMRs display TCM and TCdM (i.e. non-additive interactions) and more than 88% of them display additive methylation interaction (NI-DMRs) in the hybrid (Fig. 3B and 3D). The percentages of TCM and TCdM increased in panicle (Fig. 3B). There were 294 to 558 genes with TCM or TCdM at CG sites (Fig. 3D), while and relatively few genes with TCM or TCdM were found at non-CG sites (Fig. 3D; Supplemental Fig. S6B). However, the TCM/TCdMs showed little overlapped between shoot and panicle (Fig. 3E). The reciprocal hybrid MZ showed a similar numbers of TCM and TCdM in panicle, about 40% of which overlapped with those detected in SY63 (Fig. 3E). The analysis indicated that in the regions where the parental genomes display differential methylation additive interaction between alleles mainly takes places in the hybrid.

About 2% of the parental similarly methylated regions (SMRs) showed non-additive methylation, which however correspond to >86% of the total non-additive methylation in the hybrid (Supplemental Fig. S6B; Supplemental Fig. S7A). The remaining SMRs showed additive methylation in the hybrid. The non-additive methylation detected in the SMR principally concerns CHH and CG sequences (Supplemental Fig. S6B and S7B). However, in shoot only about 16% of non-additive CHH methylation bins were associated with 21- and 24-nt siRNAs in SY63 (Supplemental Fig. S7D). The percentage increased to 30% in panicle, which is consistent with the higher portions of 21 and 24-nt siRNA among the total siRNAs in panicle compared with shoot (Supplemental Fig. S7C and S7D). The analysis suggests that besides the RdDM pathway that is suggested to regulate non-additive methylation interaction in the hybrid (Zhang et al., 2016), additional mechanisms are likely to be involved. Analysis of expression levels of the genes with non-additive methylation revealed no difference between the hybrid and either of the parents or the mid-parent value (MPV), although a small percentage of the genes showed up and down-regulation in the hybrid (Supplemental Fig. S8).

To get information on how the parental CG and CHG DMRs are remodeled in the hybrid, we investigated allelic methylation levels in the hybrid of the SNP-bearing DMRs between ZS97 and MH63 (Supplemental Fig. S5). The analysis showed that parental CG and CHG DMRs were essentially conserved in the respective allelic bins in the hybrid (Fig. 4), indicating that parental CG and CHG methylation variations is essentially maintained in the parental alleles in the hybrid.

**Figure 4.**
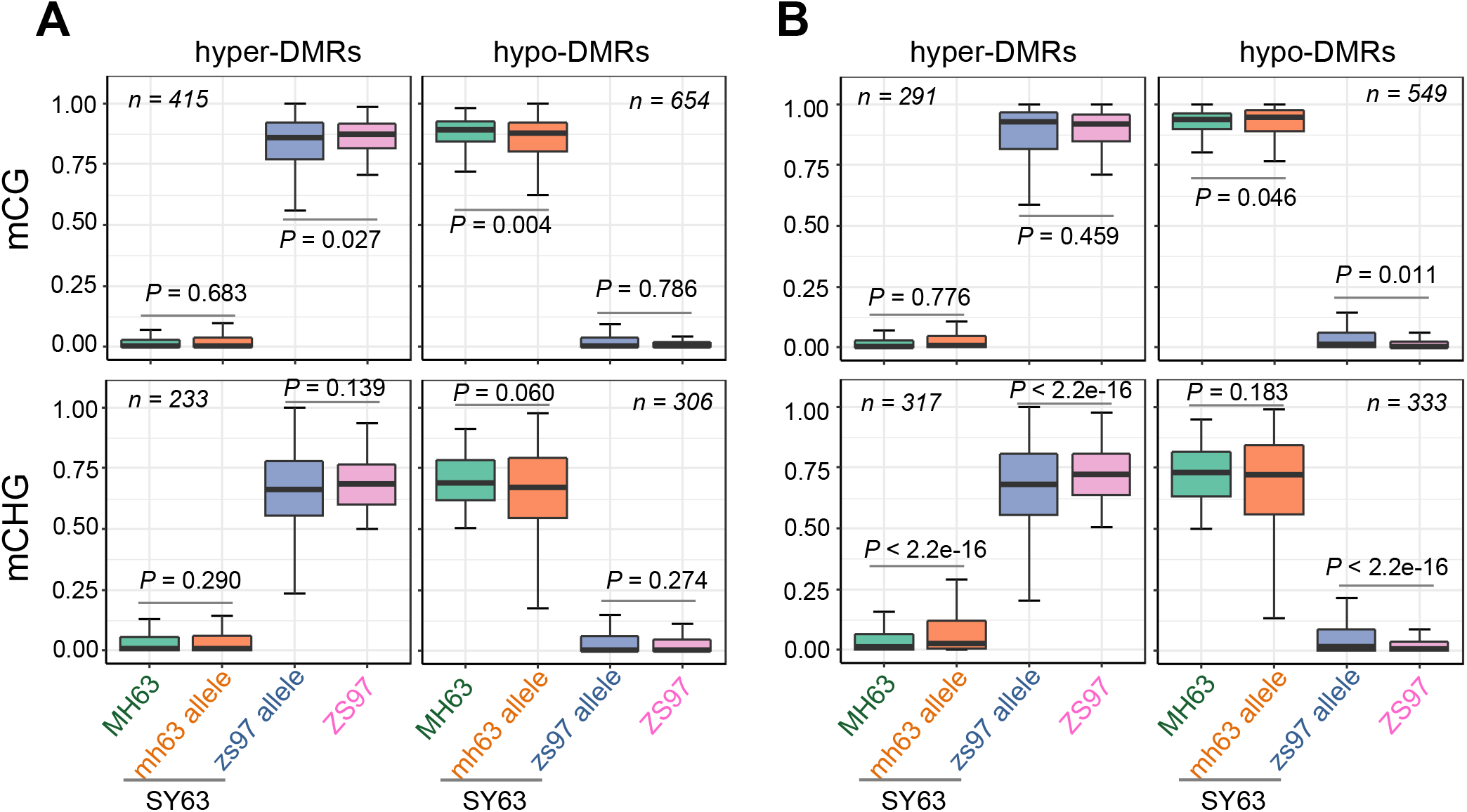
Parental methylation variations is maintained in the parental alleles in the hybrid SY63. Boxplot showing the methylation levels of DMRs (with SNPs) identified between the parental lines (MH63 and ZS97) and allele-specific methylation in hybrid (SY63) shoot (A) and panicle (B). *P* < 0.05 indicates statistical significance (Wilcoxon rank-sum test).

### Allelic-specific expression (ASE) is associated with DNA methylation

To investigate whether parental DNA methylation variations are associated with allelic-specific expression (ASE) in the hybrids, we analyzed shoot and panicle transcriptomes of the ZS97 and MH63 and their reciprocal hybrids SY63 and MZ. We used the 1,284,423 SNPs between MH63 reference sequnece2 (MH63RS2) and ZS97 reference sequence2 (ZS97RS2) identified previously for ASE calling to identify ASE genes (ASEGs) (see Methods) (Shao et al., 2019). A total of 1943 ASEGs are identified in SY63 shoot, 894 with MH63 allele-biased and 1049 with ZS97 allele-biased expression (Fig. 5A, Supplemental Dataset S3). In SY63 panicle, 533 genes showed MH63 allele-biased expression, 656 genes with ZS97 allele-biased expression (Fig. 5A). About 35-64% of the MH63 or ZS97 biased ASEG were maintained in shoot and panicle with the same biased direction (Supplemental Fig. S9A). In the reciprocal hybrid MZ shoot, the ASEG numbers were comparable with that in SY63, but in panicle higher numbers of ASEG were detected (Fig. 5A, Supplemental Dataset S4). Similarly, 38-68% of the same parent biased ASEG were persistently expressed in MZ shoot and panicle (Supplemental Fig. S9A). We noticed that most of the ASEG identified in SY63 shoot (651/894 for MH63-biaised and 755/1049 for ZS97-biaised) and panicle (518/533 for MH63-biaised and 621/656 for ZS97-biaised) were also detected in the reciprocal hybrid MZ (Supplemental Fig. S9B), suggesting that a majority of the ASEGs were independent of the parent-of-origin. Comparison of all ASEGs (a total of 4373 genes) identified in shoot and panicle of SY63 and MZ revealed 654 ASEGs that were consistently present in the reciprocal hybrids and in shoot and panicle, which includes 283 MH63-biased and 366 ZS97-biased ASEGs (Fig. 5B, Supplemental Dataset S5). The remaining 3719 genes showed inconsistent ASE directions among the reciprocal hybrids and analyzed tissues. The results were consistent with previously reported results (Shao et al., 2019). Among the identified ASEGs, many were previously shown to be associated with important agronomic traits and/or related to heterosis, such as Ghd7.1, Ghd8, MOC1, OsSPL13, OsMADS56, Xa1, etc. (Table 1) (Yan et al., 2011; Yan et al., 2013; Si et al., 2016; Shao et al., 2019). The majority (60-77%) of the ASEGs in the shoot and panicle of the reciprocal hybrids were associated with DNA methylation (Fig. 5C), which were several folds higher than the genomic average (Tan et al., 2016). Among the methylation-associated ASEG, more than two thirds showed additive methylation in the hybrids (Fig. 5C). The ASEGs showed similar difference of expression in the parental lines suggesting that ASE of the genes may be inherited from the parental expression pattern (Fig. 5D).

**Figure 5.**
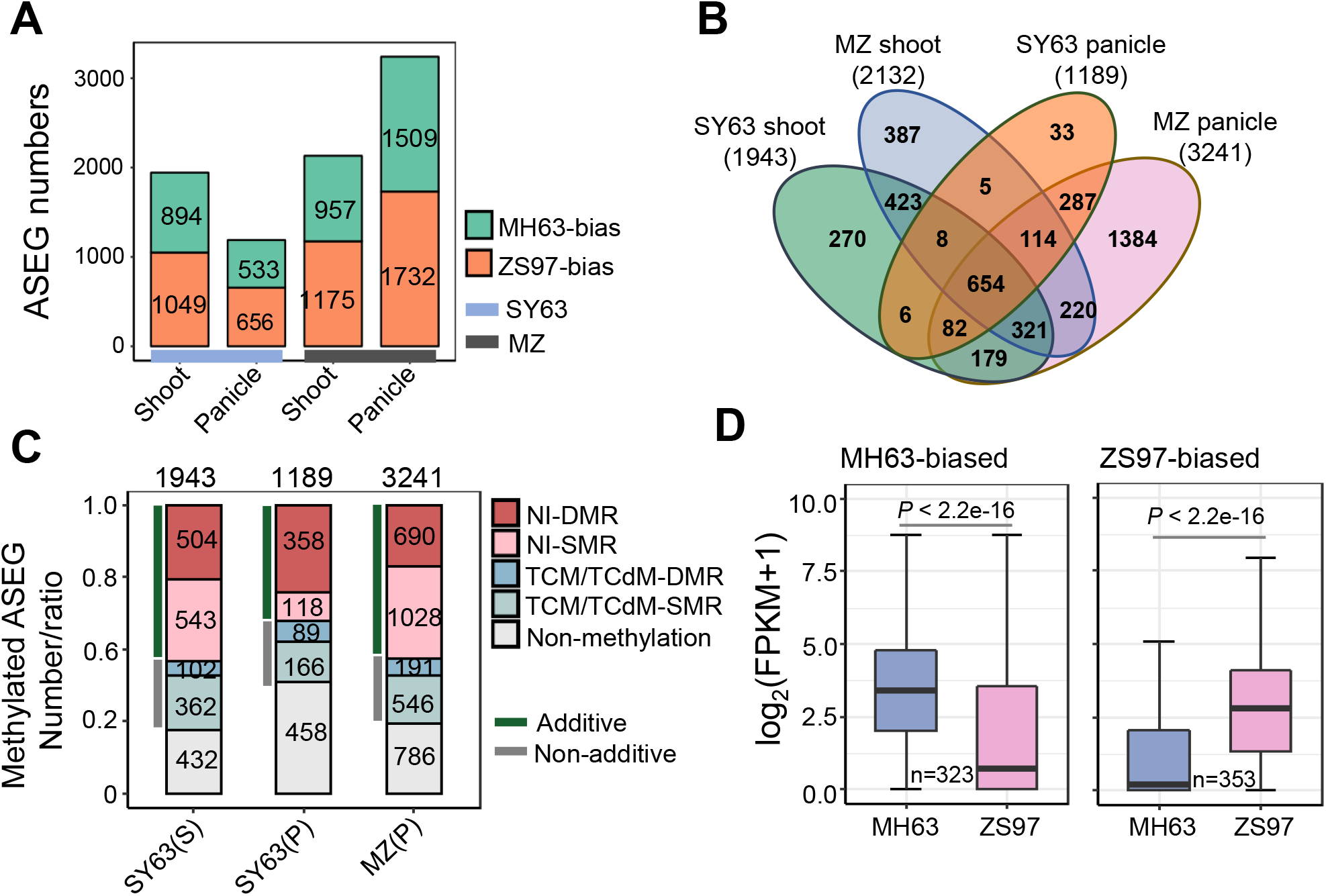
Identification of allele-specific expression genes (ASEGs) in the reciprocal hybrids shoot and panicle. (A) Numbers of ASEGs identified in SY63 and MZ shoot or panicle. (B) Venn diagram showing the numbers of consistent ASEGs in shoot and panicle of SY63 and MZ. (C) Methylated ASEG numbers or ratio in SY63 shoot or panicle and in MZ panicle. (D) Boxplot showing the difference expression between the parents of MH63-biased or ZS97-biased ASEGs identified in SY63 panicle. *P* < 0.05 indicates statistical significance (two-tailed *t* test).

**Table 1.** The previously identified important for agronomic traits or related to heterosis genes in the identified ASEGs in SY63 or MZ.

| Gene Symbol | MH63 Locus | Allele type(S) | Allele type(P) | ZS97 Locus | ASEG type |
| --- | --- | --- | --- | --- | --- |
| Ghd7.1/DTH7 | OsMH_07G0481500 | ZS97 | ZS97 | OsZS_07G0493800 | Consistent |
| OsMADS30 | OsMH_06G0437500 | ZS97 | ZS97 | OsZS_06G0417800 | Consistent |
| Xa1 | OsMH_04G0491400 | ZS97 | ZS97 | OsZS_04G0493900 | Consistent |
| Rf1a/Rf5 | OsMH_10G0314300 | ZS97 | ZS97 | OsZS_10G0336200 | Consistent |
| OsVIL1 | OsMH_08G0115500 | MH63 | MH63 | OsZS_08G0106100 | Consistent |
| Rf4,PPR782a | OsMH_10G0311800 | MH63 | MH63 | OsZS_10G0333900 | Consistent |
| MOC1 | OsMH_06G0494800 | ZS97 | NA | OsZS_06G0364100 | Inconsistent |
| LAZY1 | OsMH_11G0270000 | ZS97 | NA | OsZS_11G0283700 | Inconsistent |
| Ghd8/DTH8 | OsMH_08G0066000 | MH63 | NA | OsZS_08G0071600 | Inconsistent |
| OsMADS22 | OsMH_02G0527200 | MH63 | MH63 | OsZS_02G0620600 | Inconsistent |
| 14-3-3/GF14f | OsMH_03G0497400 | MH63 | NA | OsZS_03G0494600 | Inconsistent |
| Wx | OsMH_06G0027500 | MH63 | NA | OsZS_06G0028900 | Inconsistent |
| Xa26 | OsMH_11G0447800 | MH63 | MH63 | OsZS_11G0447800 | Inconsistent |
| TAW1 | OsMH_10G0384000 | MH63 | NA | OsZS_10G0317200 | Inconsistent |
| OsGID1 | OsMH_05G0309000 | NA | MH63 | OsZS_05G0316500 | Inconsistent |
| OsSPL13 | OsMH_07G0316100 | NA | MH63 | OsZS_07G0327600 | Inconsistent |
| OsYUCCA3 | OsMH_01G0511100 | NA | MH63 | OsZS_01G0717200 | Inconsistent |
| OsGA2OX3 | OsMH_01G0531100 | NA | MH63 | OsZS_01G0526400 | Inconsistent |
| Hd1 | OsMH_06G0155400 | NA | MH63 | OsZS_06G0152400 | Inconsistent |
| OsMADS15 | OsMH_07G0007900 | NA | MH63 | OsZS_07G0006000 | Inconsistent |

(S, shoot; P, panicle)

### Allelic-specific CHG methylation is associated with allelic gene expression

To study whether the methylated ASEG is associated with allelic-specific methylation, we analyzed methylation levels of ASEGs (gene body and 2 kb 5’ and 3’ flanking regions) in the parents and the hybrid parental alleles at CG, CHG and CHH contexts. For the 533 panicle MH63-biased ASEGs, 322 to 336 have CG, CHG, and/or CHH methylation and for the 656 panicle ZS97-biaised ASEGs, 377 to 395 have CG, CHG, and/or CHH methylation (Fig. 6A, Supplemental Dataset S6). Metaplot analysis revealed that MH63-biased ASEGs showed lower gene body CHG methylation in the MH63 alleles than the ZS97 alleles in the hybrid (Fig. 6A). Conversely, in the 384 (out of 656) ZS97-biased ASEGs that show CHG methylation (Supplemental Dataset S6) the ZS97 alleles display lower CHG methylation than the MH63 alleles in the hybrid (Fig. 6A). Quantitative analysis indicated that the body region of the ZS97 alleles showed significantly higher CHG methylation than the MH63 alleles in the MH63-biaised ASEGs and lower CHG methylation in the ZS97-biaised ASEGs (Fig. 6B), indicating that allele-specific methylation (ASM) at CHG sites was negatively correlated with ASE, consistent with the repressive role of CHG methylation in gene expression. This pattern of ASM was detected in several agronomically important genes such as MOC1, GME, Xa26, and MADS15 (Fig. 6C and 6D, Supplemental Fig. S10, Supplemental Dataset S6) (Liao et al., 2019; Shao et al., 2019). At the CG context, the MH63-biased ASEGs showed higher gene body methylation in the MH63 alleles than the ZS97 alleles, indicating that gene body CG ASM positively correlated with ASE, corroborating the positive effect of gene body CG methylation for gene expression (Bewick and Schmitz, 2017). However, there was no discernable difference of allelic-specific CG methylation of the ZS97-biased ASEGs in the hybrid. This might be due to overall lower CG methylation in the ZS97 alleles compared with the MH63 alleles of the ASEGs (Fig. 6B). At CHH context, no clear ASM could be discerned, except that methylation peaks could be observed mainly in the flanking regions in both alleles of MH63-biased or ZS97-biased ASEGs. Similar trends were observed for the SY63 shoot ASEGs and the reciprocal hybrid (MZ) panicle ASEGs (Supplemental Fig. S11 and S12; Supplemental Dataset S7 and S8). However, the ASM differences in SY63 shoot or MZ panicle were weaker, probably because larger numbers of ASEGs were taken into consideration. Taken together, the analysis identifies a role of ASM at gene body CHG and, to a lesser extent, CG sites in ASE.

**Figure 6.**
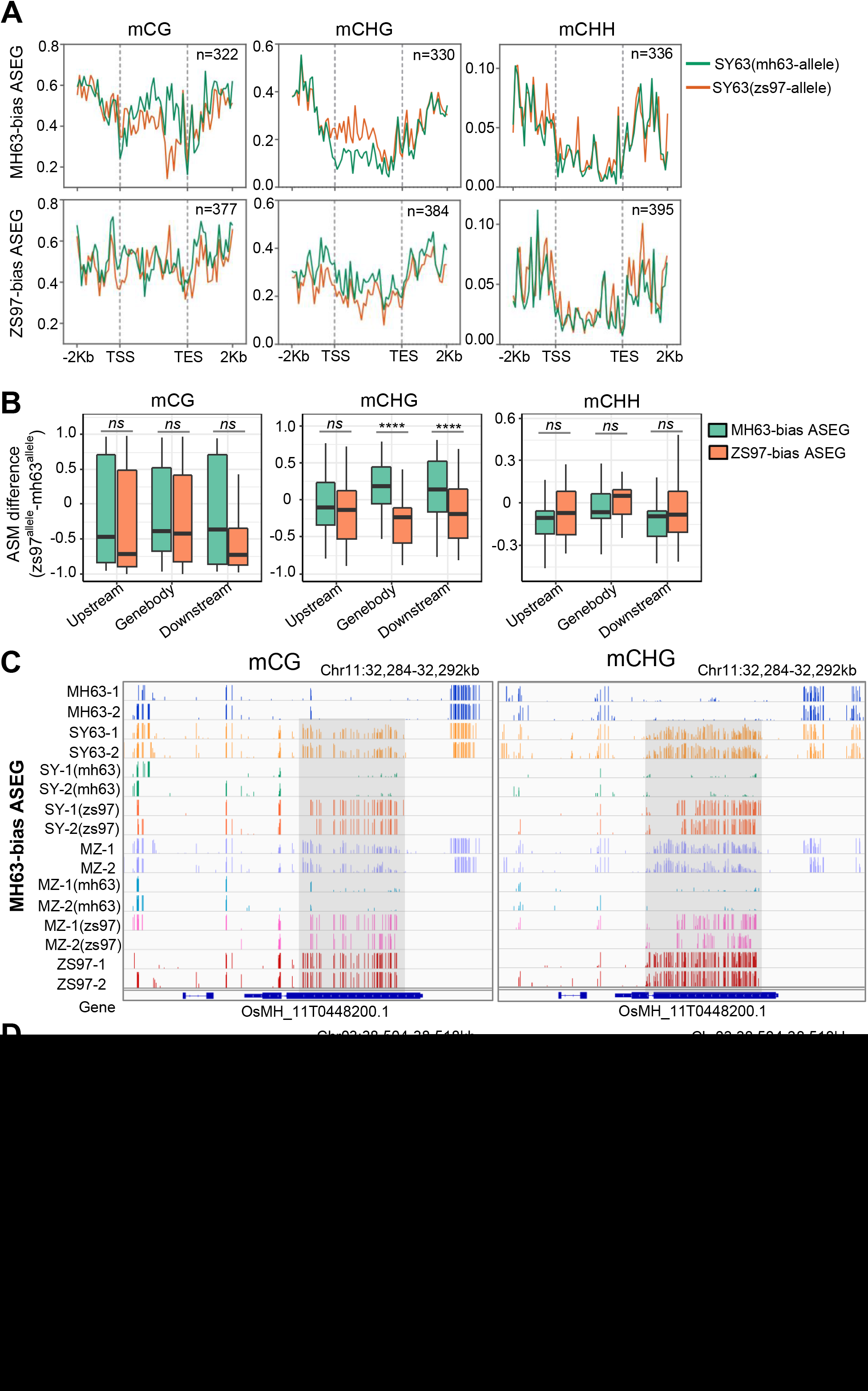
Differential allele-specific methylation of the ASEGs identified in the hybrid SY63 panicle. (A) Metaplot showing CG or CHG methylation difference between MH63-alleles and ZS97 alleles of MH63-biased or ZS97-biased ASEG in SY63 panicle. (B) Boxplot showing allele-specific methylation (ASM) differences (ZS97 allele-MH63 allele) of MH63-biased ASEG and ZS97-biased ASEG in SY63 panicle. ****, *P* < 0.0001; *ns*, *P* > 0.05 (Wilcoxon rank-sum test). (C) Genome browser screenshots showing allelic-specific CG or CHG methylation of a MH63-biased ASEG (Xa26, OsMH_11T0448200) in the reciprocal hybrids SY63 and MZ panicle in comparison with the hybrid (average) and the parental methylation levels. Two biological replicates are shown. SY (mh63 or zs97) and MZ (mh63 or zs97) indicate the MH63 or ZS97 allele in SY63 or MZ. (D) Genome browser screenshots showing allelic-specific CG or CHG methylation of a ZS97-biased ASEG (OsMH_03T0611800) between the parents allele in the hybrids panicle.

## DISCUSSION

### DNA methylation divergence between the parental lines ZS97 and MH63

This work reveals more important methylation variations at CG and CHG than CHH sites between the parental lines of the elite hybrid rice SY63, although the CHH methylation variation increases in panicle. This is unexpected, as CHH methylation appears to be more stochastic and tends to be more variable in population and during development (Higo et al., 2020) (as shown in Fig. 1D). Despite that CG and CHG methylation mainly target TEs and repetitive sequences in the rice genome (Tan et al., 2016) a majority of the CG and CHG DMRs between ZS97 and MH63 are located in genes (Fig. 2B). Since only the syntenic regions (with or without SNP, but excluding InDels) between the parental genomes were taken into consideration, it is unlikely that the CG and CHG methylation variations are solely resulted from the parental DNA sequence divergence, albeit higher percentages of DMRs detected in SNP bins (Supplemental Fig. S5). In addition, the work reveals difference of overall methylation levels and variation during development between MH63 than ZS97. Whether the degree of methylation difference between the parental lines is related to hybrid vigor and can be used as a criterion to allow predicting heterotic performance awaits further investigation.

The difference in gene body CG and CHG methylation between MH63 and ZS97 may have arisen during their breeding and adaptation in different subcontinental regions. It has been shown that gene body CG methylation is present only in plant species that contain CMT3 that catalyzes CHG methylation, but absent from those without CMT3 (Bewick et al., 2016). Recent data showed that expressing *CMT3* in a species naturally lacking CG body methylation reconstitutes gene body CG methylation through a progression of *de novo* CHG methylation on expressed genes, followed by the accumulation of CG methylation that could be inherited even following loss of the *CMT3* transgene (Wendte et al., 2019). This would suggest that differential CMT3 activities between the parental lines could be related to the CG and CHG methylation divergence. However, no difference of CMT3 amino acid sequence is detected between ZS97 and MH63 (Supplemental Fig. S2C). By contrast, CMT2 that is involved in both CHG and CHH methylation presents a large deletion in ZS97 (Supplemental Fig. S2C). Whether the CMT2 polymorphism is related to the difference in CG and CHG methylation await further analysis. Alternatively, difference in chromatin modification such as H3K9me2 may also contribute to the difference in CG and CHG methylation between the two parental lines.

### Interaction of parental DNA methylation epigenomes in hybrid rice

DNA methylation remodeling in hybrid leading to variation relative to the parental epigenomes has been proposed to be involved in heterosis (Greaves et al., 2012; Shen et al., 2012; Zhang et al., 2016). Processes such as the non-additive interactions between the parental alleles leading to either gain (TCM) or loss (TCdM) of allelic methylation have been found to contribute to the observed methylation changes in hybrids (Greaves et al., 2012; Shivaprasad et al., 2012; Groszmann et al., 2013; Chen et al., 2014; Zhang et al., 2016). The present data show that in the hybrid SY63, only a small percentage of bins show non-additive methylation, most of which is detected in the genomic regions where parental DNA methylations are similar (Supplemental Fig. S6 and S7B). Due to the presence of siRNAs at regions that undergo TCM/TCdM, it has been suggested that siRNAs are the initiating molecules that establish these TCM/TCdM events (Greaves et al., 2015). However, in the rice hybrids, only about 15% in shoot and 30% in panicle of the non-additive (both hyper and hypo) DNA methylation regions are associated with 21- and 24-nt siRNA (Supplemental Fig. S7D). In addition, besides CHH methylation, non-additive interaction also involves CG methylation in SY63. Together, the observations suggest that besides RdDM that is suggested to be important for communication between alleles (Zhang et al., 2016), additional mechanisms are likely to be involved in the non-additive methylation interaction between alleles in the hybrid rice. That no overall expression difference of the non-additive methylation genes is observed between the hybrid and either of the parents or the mid-parent values (Supplemental Fig. S8) supports the hypothesis that non-additive methylation variation may principally have impact at specific loci rather than on a global scale.

### Allelic-specific methylation (ASM) at CHG sites represses Allelic-specific expression (ASE)

It is suggested that ASE can lead to phenotypic variation and play a role in heterosis (Springer and Stupar, 2007; Paschold et al., 2012; Goff and Zhang, 2013; Shao et al., 2019). This notion is supported by the identification of several ASE genes involved in key agronomic traits and or hybrid vigor in this and previous study (Supplemental Dataset S3, S5, Table 1) (Shao et al., 2019). Although genetic variations may cause gene expression difference between alleles, it has been shown that epigenetic mechanism is involved in ASE. In mammals, ASE is often associated with epigenetic inactivation in X-chromosome and genomic imprinting as well as non-imprinted genes (Knight, 2004). In plants, hundreds of imprinted genes in endosperm or embryo display allelic differences (effect of the parent-of-origin) in DNA methylation and chromatin modification (such as the histone modification mark H3K27me3), which are important for establishing and maintaining imprinted expression (Gehring, 2013). It appears that the paternal allele of paternally expressed genes in endosperm of several plant species is associated with DNA methylation, and the silent maternal allele shows hypo DNA methylation and is marked by H3K27me3 (Zhang et al., 2014; Moreno-Romero et al., 2016). Unlike the imprinted genes, nearly all of the ASEG in the SY63 are detected in the reciprocal hybrid MZ (Supplemental Fig. S9B), thus show no effect of the parent-of-origin. The observation that most of the identified ASEGs in the rice hybrids are methylated suggests DNA methylation may be involved in the ASE. The reverse correlation between ASE and allelic-specific methylation (ASM) at CHG sites suggests that CHG methylation maintains or reinforces repression of the parental alleles in the hybrids, consistent with the observation that genes with variation in CHG methylation show difference of expression between the parental lines (Fig. 2E). This is reminiscent of the finding that the silent maternal allele of paternally expressed genes in endosperm of *Arabidopsis lyrata* is marked by hyper CHG methylation (Klosinska et al., 2016). Increase of maternal allele CHG methylation was associated with higher expression bias in favor of the paternal allele (Klosinska et al., 2016). The present results indicate that allelic-specific gene body CHG methylation is likely inherited from the parental epigenomes and is maintained or reinforced in a allelic-specific manner in the hybrid and during development (Fig. 4, Supplemental Fig. S10), possibly though the reinforcing feedback loop with H3K9me2 (Law and Jacobsen, 2010). As nearly all of the genes with ASMs at CHG also show CG methylation which, by contrast, positively correlates with ASE, retention of allelic-specific CG gene body methylation could possibly protect them from gain of CHG methylation. This is consistent with previous results showing that in hybrids of Arabidopsis *Ler* and C24 ecotypes both cis- and trans-regulated DNA methylation play roles in ASM, with cis-regulation playing a major role in CG methylation and trans-regulation (e.g. involving siRNA, DNA methylation regulators, etc.) playing major roles in CHG and CHH methylation (Chen et al., 2010). It was shown that in Arabidopsis DDM1 mutation reduces heterosis level of biomass (Kawanabe et al., 2016; Zhang et al., 2016), while mutations in RNA polymerase IV (involved in siRNA production and RdDM) or MET1 (required for CG methylation) had no effect on heterosis (Kawanabe et al., 2016). As DDM1 is required mainly for CG and CHG methylation in both Arabidopsis and rice (Zemach et al., 2013; Tan et al., 2016; Tan et al., 2018), the effect of DDM1 mutation on heterosis is in favor of the hypothesis that CHG methylation variation may play a role in epigenetic mechanism involved in heterosis.

## MATERIALS AND METHODS

### Plant materials and growth conditions

The *Indica/Xian* rice varieties Zhenshan 97 (ZS97) and Minghui 63 (MH63) and their reciprocal hybrids [Shanyou 63 (SY63) and MH63/ZS97 (MZ)] were used in this study. Germinated seeds were planted under growth chambers with condition of 14h light and 10h darkness and with temperature of 32°C/28°C. Seedling shoot at 4-leaf stage were collected for total RNA or genomic DNA extraction. The plants used for collection of young panicle were grown in field and managed under normal agricultural conditions on the Huazhong Agricultural University rice experimental field, Wuhan, China. Young panicle with 2-mm length at the early stage of floral organs differentiation were collected and immediately placed in liquid N2 and stored at −80°C until RNA and genomic DNA extraction.

### RNA-seq data analysis and ASE identification

Total RNA was extracted from the indicated tissues using TRIzol reagent (Invitrogen, USA) according to the manufacturer’s protocol. RNA concentrations were measured using Nanodrop 2000 (Thermo Scientific, USA) and further quantified using Qubit 2.0 Fluorometer (Invitrogen, USA). A total of 2 μg RNA was used for mRNA isolated and RNA-seq library construction using the TruSeq RNA Library Preparation Kit (Illumina, USA), according to the manufacturer’s recommendations. The libraries were sequenced using Illumina HiSeq 2000 platform.

RNA-seq raw reads were filtered to remove adapter and low-quality bases by Trimmomatic (version 0.35). Allele-specific-expression (ASE) reads separation and identification were carried out as previously described (Shao et al., 2019). Briefly, the cleaned reads of 24 libraries (4 genotypes x 2 tissues x3 replicates) were aligned to two parental reference genomes (MH63RS2 and ZS97RS2 from Rice Information GateWay [RIGW], http://rice.hzau.edu.cn/rice/) respectively by HISAT2 (version 2.1.0) using default parameters (http://ccb.jhu.edu/software/hisat2/index.shtml). For the mapping reads, a stringent filtration was conducted using a customized perl script. It was retained when the filtered reads can be perfectly mapped to one parental genome or with SNPs mapped to the other parental genome. The trimmed high-quality reads from the hybrid were divided into 6 sets, according to the reads of each replicate aligned to MH63RS2 and ZS97RS2, to separate MH63- or ZS97-specific reads in the SNP calling. A total of 1,284,423 SNPs between MH63RS2 and ZS97RS2 were used for ASE calling as the reference. The identification of ASE for each gene was based on the SNPs between two parental genomes, and it needs to meet the following three criteria: (1) every SNP is covered by no less than 5 reads, (2) the ratio of read counts of SNPs from the two parental alleles differs significantly from 1:1 (Q < 0.01), and (3) the significant bias of different SNPs of the same gene is not in different directions. A gene showing ASE no less than one SNP is referred to as an allele-specific-expression gene (ASEG).

Gene expression levels were calculated by StringTie (version1.2.1) with parameters for strand-specific RNA-seq (Pertea et al., 2016). Differentially expressed gene of statistical significance (adjust P < 0.01 and abs [log_2_ (fold change)] > 1) were identified between ZS97 and MH63 by the nbinomTest function of the DESeq package (Anders and Huber, 2010).

### Bisulfite-seq library construction and WGBS-seq data analysis

Genomic DNA was extracted using DNeasy plant mini kit (QIAGEN, Germany) according to the manufacturer’s instructions. Bisulfite conversion of DNA, library construction and sequencing were performed at the Beijing Genomics Institute.

Sequence quality of the whole-genome bisulfite sequencing (WGBS) data was evaluated by FastQC (version 0.11.5). The adapter and low-quality reads were removed by Trimmomatic (version 0.35), and the clean data were mapped to the MH63RS2 reference genome by Bismark (version 0.22.3) using default parameters (Krueger and Andrews, 2011). The uniquely mapped reads were retained for further analysis. Individual cytosines with more than four reads were considered for DNA methylation level calculation.

Kernel density plots were generated by comparing the average cytosine methylation level within 200-bp window between two samples, the window at least contained five cytosines and every cytosine that were covered by at least five reads were used.

### Identification of DMRs and Analysis of methylation interaction in F1 hybrids

To identify the differentially methylated regions (DMRs), the whole genome was divided into 200-bp bins. Bins that contained at least five cytosines each and every cytosine with at least a five-fold coverage were retained. Bins with absolute methylation difference of 0.7, 0.5, 0.2 for CG, CHG and CHH sites respectively, and *P*-value>0.01 (Fisher’s exact test) between comparisons was considered as DMRs. For the methylation difference between MH63 and ZS97, excluded from the identified DMRs, the remaining bins were considered as similarly methylated regions (SMRs).

Identification of methylation interaction in F1 hybrid was conducted according to a previous study (Schultz et al., 2012; Zhang et al., 2016). The weighted methylation level of F1 hybrid for each DMR between MH63 and ZS97 was calculated as the predicted additive value (PAV). In brief, the combined parental methylated reads were divided by the combined parental total reads inside parental DMR by the following formula:

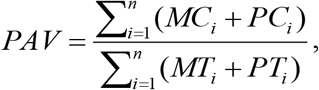

i = position of cytosine, n = total number of cytosine positions in the DMR, MC = maternal methylated reads, PC = paternal methylated reads, MT = maternal methylated and unmethylated reads, PT = paternal methylated and unmethylated reads.

To compare the methylation level in F1 hybrids with parental DMRs, Fisher’s exact test was used to validate the methylation difference between F1 hybrid and corresponding PAV. The FDR was generated from an adjusted *P*-value using the Benjamini-Hochberg method. The F1 hybrid bins whose methylation levels were higher than the corresponding PAV with FDR < 0.01 were considered as trans-chromosomal methylation regions (TCM-DMRs), whereas bins with F1 methylation levels were lower than the PAV and FDR < 0.01 were considered as trans-chromosomal demethylation regions (TCdM-DMRs), the remaining bins with FDR > 0.01 were considered as non-interaction methylation regions (NI-DMRs). The comparison of methylation difference between F1 hybrid and parental SMRs, the calculation method was the same as described above.

### sRNA-seq data analysis

Analysis of small RNA-seq data was performed as previously described (Tan et al., 2016). Briefly, sRNA-seq raw data were cleaned by removing adaptor and low-quality reads by Trimmomatic (version 0.35). MicroRNA (http://www.mirbase.org/ftp.shtml), pre-miRNA, tRNA and rRNA were removed. The cleaned reads were aligned to the MH63RS2 reference genome by Bowtie (version 1.3.0, https://sourceforge.net/projects/bowtie-bio/files/bowtie/1.3.0/), allowing zero mismatches and unique mapping. Two types of small RNA abundance were calculated by 21- or 24-nucleotide read numbers falling into 1-kb windows/regions throughout the whole genomes.

### Allele-specific methylation (ASM) analysis

To separate the MH63-allele or ZS97-allele reads from the hybrids BS-seq data, SNPs in the syntenic block regions of the parental genomes were used to filtrate the BS-seq reads. A total of 1,284,423 SNPs between two parents were identified by comparison the high quality genome sequences of MH63 and ZS97 using Mummer3.23 (https://sourceforge.net/projects/mummer/files/). Firstly, the filtered reads of BS-seq libraries were aligned to the parental reference genomes (MH63RS2 and ZS97RS2) by Bismark (version 0.22.3) using default parameters. Then the mapped reads of F1 hybrids were separated into MH63-allele or ZS97-allele reads, according to the identified SNPs between MH63RS2 and ZS97RS2 and its location in the synteny block using a customized perl script. To assess the accuracy of the separated reads, the parental reads were also mapped to two reference genomes by the same process. Finally, the allele-specific-methylation level of hybrid F1 were calculated based on the separated MH63-allele or ZS97-allele reads independently by BatMeth2 (version 2.01) using default parameters (Zhou et al., 2019). Fisher’s exact tests were used to examine the methylation difference between MH63-allele and ZS97-allele reads within 200-bp windows.

### Accession numbers

The BS-seq and RNA-seq data produced in this study are deposited to NCBI Sequence Read Archive (BioProject ID: PRJNA664649) (https://submit.ncbi.nlm.nih.gov/subs/sra/SUB8163360/overview).

## Supplemental Data

**Supplemental Figure S1.** BS-seq data and DNA methylation levels of MH63, ZS97 and their reciprocal hybrids.

**Supplemental Figure S2.** Expression levels and amino acid sequence comparison of DNA methylation maintenance genes in rice.

**Supplemental Figure S3.** Heatmap and Pearson correlation coefficients of RNA-seq replicates of MH63 (MH), ZS97 (ZS), SY63 (SY) and MH63/ZS97 (MZ) in shoot (S) and panicle (P).

**Supplemental Figure S4.** Genome browser screenshots of DNA methylation in MH63 and ZS97 shoot or panicle.

**Supplemental Figure S5.** DMRs with SNPs or without SNPs between MH63 and ZS97 in shoot and panicle.

**Supplemental Figure S6**. Identification of methylation interaction in hybrids panicle and shoot.

**Supplemental Figure S7.** Methylation interaction in hybrids panicle and shoot.

**Supplemental Figure S8.** The expression levels of genes with SMR-TCM or SMR-TCdM in SY63 panicle.

**Supplemental Figure S9.** Identification of allele-specific expression genes (ASEGs) in the hybrids shoot and panicle.

**Supplemental Figure S10.** Genome browser screenshots of allele-specific methylated ASEGs.

**Supplemental Figure S11.** Differential allele-specific methylation of the ASEGs identified in SY63 shoot.

**Supplemental Figure S12.** Differential allele-specific methylation of the ASEGs identified in MZ panicle.

**Supplemental Table S1.** Information of BS-seq data of MH63, ZS97 and their reciprocal hybrids F1 in shoot and panicle.

**Supplemental Table S2.** Information of RNA-seq data of MH63, ZS97 and their reciprocal hybrids F1 in shoot and panicle.

**Supplemental Dataset S1.** Differentially methylated genes identified between ZS97 and MH63 in CG, CHG and CHH context in panicle.

**Supplemental Dataset S2.** Differentially methylated genes identified between ZS97 and MH63 in CG, CHG and CHH context in shoot.

**Supplemental Dataset S3.** Information of the ASEGs identified in SY63 shoot and panicle.

**Supplemental Dataset S4.** Information of the ASEGs identified in MZ shoot and panicle.

**Supplemental Dataset S5.** Information of the consistent ASEGs identified in hybrid F1.

**Supplemental Dataset S6.** Allele-specific methylated ASEGs of MH63-biased or ZS97-biased identified in SY63 panicle.

**Supplemental Dataset S7.** Allele-specific methylated ASEGs of MH63-biased or ZS97-biased identified in SY63 shoot.

**Supplemental Dataset S8.** Allele-specific methylated ASEGs of MH63-biased or ZS97-biased identified in MZ panicle.

## ACKNOWLEDGMENTS

We thank Drs Xianghua LI and Jinghua Xiao for assistance.

## Author Contributions

X.M. did most of the experimental work and data analysis; F.X. did bioinformatics analysis; Q.J. and Q.Z. did field work; T.H., B.W., L.S., and Y.Z. participated in experimental work, Q.Z. and D.-X.Z. designed and supervised the work; D.-X.Z. wrote the paper with inputs from X.M.

## LITERATURE CITED

Anders S, Huber W (2010) Differential expression analysis for sequence count data. Genome Biol 11: R106

Barber WT, Zhang W, Win H, Varala KK, Dorweiler JE, Hudson ME, Moose SP (2012) Repeat associated small RNAs vary among parents and following hybridization in maize. Proc Natl Acad Sci U S A 109: 10444–10449

Bewick AJ, Ji LX, Niederhuth CE, Willing EM, Hofmeister BT, Shi XL, Wang L, Lu ZF, Rohr NA, Hartwig B, Kiefer C, Deal RB, Schmutz J, Grimwood J, Stroud H, Jacobsen SE, Schneeberger K, Zhang XY, Schmitz RJ (2016) On the origin and evolutionary consequences of gene body DNA methylation. Proceedings of the National Academy of Sciences of the United States of America 113: 9111–9116

Bewick AJ, Schmitz RJ (2017) Gene body DNA methylation in plants. Curr Opin Plant Biol 36: 103–110

Chen F, He G, He H, Chen W, Zhu X, Liang M, Chen L, Deng XW (2010) Expression analysis of miRNAs and highly-expressed small RNAs in two rice subspecies and their reciprocal hybrids. J Integr Plant Biol 52: 971–980

Chen S, He H, Deng X (2014) Allele-specific DNA methylation analyses associated with siRNAs in Arabidopsis hybrids. Sci China Life Sci 57: 519–525

Chen ZJ (2013) Genomic and epigenetic insights into the molecular bases of heterosis. Nat Rev Genet 14: 471–482

Chodavarapu RK, Feng S, Ding B, Simon SA, Lopez D, Jia Y, Wang GL, Meyers BC, Jacobsen SE, Pellegrini M (2012) Transcriptome and methylome interactions in rice hybrids. Proc Natl Acad Sci U S A 109: 12040–12045

Crisp PA, Hammond R, Zhou P, Vaillancourt B, Lipzen A, Daum C, Barry K, de Leon N, Buell CR, Kaeppler SM, Meyers BC, Hirsch CN, Springer NM (2020) Variation and Inheritance of Small RNAs in Maize Inbreds and F1 Hybrids. Plant Physiol 182: 318–331

Dapp M, Reinders J, Bediee A, Balsera C, Bucher E, Theiler G, Granier C, Paszkowski J (2015) Heterosis and inbreeding depression of epigenetic Arabidopsis hybrids. Nat Plants 1: 15092

Gehring M (2013) Genomic imprinting: insights from plants. Annu Rev Genet 47: 187–208

Goff SA, Zhang Q (2013) Heterosis in elite hybrid rice: speculation on the genetic and biochemical mechanisms. Curr Opin Plant Biol 16: 221–227

Greaves IK, Gonzalez-Bayon R, Wang L, Zhu A, Liu PC, Groszmann M, Peacock WJ, Dennis ES (2015) Epigenetic Changes in Hybrids. Plant Physiol 168: 1197–1205

Greaves IK, Groszmann M, Wang A, Peacock WJ, Dennis ES (2014) Inheritance of Trans Chromosomal Methylation patterns from Arabidopsis F1 hybrids. Proc Natl Acad Sci U S A 111: 2017–2022

Greaves IK, Groszmann M, Ying H, Taylor JM, Peacock WJ, Dennis ES (2012) Trans chromosomal methylation in Arabidopsis hybrids. Proc Natl Acad Sci U S A 109: 3570–3575

Groszmann M, Greaves IK, Albertyn ZI, Scofield GN, Peacock WJ, Dennis ES (2011) Changes in 24-nt siRNA levels in Arabidopsis hybrids suggest an epigenetic contribution to hybrid vigor. Proc Natl Acad Sci U S A 108: 2617–2622

Groszmann M, Greaves IK, Fujimoto R, Peacock WJ, Dennis ES (2013) The role of epigenetics in hybrid vigour. Trends Genet 29: 684–690

He G, Chen B, Wang X, Li X, Li J, He H, Yang M, Lu L, Qi Y, Wang X, Deng XW (2013) Conservation and divergence of transcriptomic and epigenomic variation in maize hybrids. Genome Biol 14: R57

He G, Zhu X, Elling AA, Chen L, Wang X, Guo L, Liang M, He H, Zhang H, Chen F, Qi Y, Chen R, Deng XW (2010) Global epigenetic and transcriptional trends among two rice subspecies and their reciprocal hybrids. Plant Cell 22: 17–33

Higo A, Saihara N, Miura F, Higashi Y, Yamada M, Tamaki S, Ito T, Tarutani Y, Sakamoto T, Fujiwara M, Kurata T, Fukao Y, Moritoh S, Terada R, Kinoshita T, Ito T, Kakutani T, Shimamoto K, Tsuji H (2020) DNA methylation is reconFigured at the onset of reproduction in rice shoot apical meristem. Nat Commun 11: 4079

Hua J, Xing Y, Wu W, Xu C, Sun X, Yu S, Zhang Q (2003) Single-locus heterotic effects and dominance by dominance interactions can adequately explain the genetic basis of heterosis in an elite rice hybrid. Proc Natl Acad Sci U S A 100: 2574–2579

Huang X, Wei X, Sang T, Zhao Q, Feng Q, Zhao Y, Li C, Zhu C, Lu T, Zhang Z, Li M, Fan D, Guo Y, Wang A, Wang L, Deng L, Li W, Lu Y, Weng Q, Liu K, Huang T, Zhou T, Jing Y, Li W, Lin Z, Buckler ES, Qian Q, Zhang QF, Li J, Han B (2010) Genome-wide association studies of 14 agronomic traits in rice landraces. Nat Genet 42: 961–967

Huang X, Yang S, Gong J, Zhao Y, Feng Q, Gong H, Li W, Zhan Q, Cheng B, Xia J, Chen N, Hao Z, Liu K, Zhu C, Huang T, Zhao Q, Zhang L, Fan D, Zhou C, Lu Y, Weng Q, Wang ZX, Li J, Han B (2015) Genomic analysis of hybrid rice varieties reveals numerous superior alleles that contribute to heterosis. Nat Commun 6: 6258

Kawanabe T, Ishikura S, Miyaji N, Sasaki T, Wu LM, Itabashi E, Takada S, Shimizu M, Takasaki-Yasuda T, Osabe K, Peacock WJ, Dennis ES, Fujimoto R (2016) Role of DNA methylation in hybrid vigor in Arabidopsis thaliana. Proc Natl Acad Sci U S A 113: E6704–E6711

Klosinska M, Picard CL, Gehring M (2016) Conserved imprinting associated with unique epigenetic signatures in the Arabidopsis genus. Nat Plants 2: 16145

Knight JC (2004) Allele-specific gene expression uncovered. Trends Genet 20: 113–116

Krueger F, Andrews SR (2011) Bismark: a flexible aligner and methylation caller for Bisulfite-Seq applications. Bioinformatics 27: 1571–1572

Lauss K, Wardenaar R, Oka R, van Hulten MHA, Guryev V, Keurentjes JJB, Stam M, Johannes F (2018) Parental DNA Methylation States Are Associated with Heterosis in Epigenetic Hybrids. Plant Physiol 176: 1627–1645

Law JA, Jacobsen SE (2010) Establishing, maintaining and modifying DNA methylation patterns in plants and animals. Nature Reviews Genetics 11: 204–220

Liao Z, Yu H, Duan J, Yuan K, Yu C, Meng X, Kou L, Chen M, Jing Y, Liu G, Smith SM, Li J (2019) SLR1 inhibits MOC1 degradation to coordinate tiller number and plant height in rice. Nat Commun 10: 2738

Moreno-Romero J, Jiang H, Santos-Gonzalez J, Kohler C (2016) Parental epigenetic asymmetry of PRC2-mediated histone modifications in the Arabidopsis endosperm. EMBO J 35: 1298–1311

Paschold A, Jia Y, Marcon C, Lund S, Larson NB, Yeh CT, Ossowski S, Lanz C, Nettleton D, Schnable PS, Hochholdinger F (2012) Complementation contributes to transcriptome complexity in maize (Zea mays L.) hybrids relative to their inbred parents. Genome Res 22: 2445–2454

Pertea M, Kim D, Pertea GM, Leek JT, Salzberg SL (2016) Transcript-level expression analysis of RNA-seq experiments with HISAT, StringTie and Ballgown. Nat Protoc 11: 1650–1667

Schnable PS, Springer NM (2013) Progress toward understanding heterosis in crop plants. Annu Rev Plant Biol 64: 71–88

Schultz MD, Schmitz RJ, Ecker JR (2012) ‘Leveling’ the playing field for analyses of single-base resolution DNA methylomes. Trends Genet 28: 583–585

Seifert F, Thiemann A, Grant-Downton R, Edelmann S, Rybka D, Schrag TA, Frisch M, Dickinson HG, Melchinger AE, Scholten S (2018) Parental Expression Variation of Small RNAs Is Negatively Correlated with Grain Yield Heterosis in a Maize Breeding Population. Front Plant Sci 9: 13

Shao GN, Lu ZF, Xiong JS, Wang B, Jing YH, Meng XB, Liu GF, Ma HY, Liang Y, Chen F, Wang YH, Li JY, Yu H (2019) Tiller Bud Formation Regulators MOC1 and MOC3 Cooperatively Promote Tiller Bud Outgrowth by Activating FON1 Expression in Rice. Molecular Plant 12: 1090–1102

Shao L, Xing F, Xu C, Zhang Q, Che J, Wang X, Song J, Li X, Xiao J, Chen LL, Ouyang Y, Zhang Q (2019) Patterns of genome-wide allele-specific expression in hybrid rice and the implications on the genetic basis of heterosis. Proc Natl Acad Sci U S A 116: 5653–5658

Shen H, He H, Li J, Chen W, Wang X, Guo L, Peng Z, He G, Zhong S, Qi Y, Terzaghi W, Deng XW (2012) Genome-wide analysis of DNA methylation and gene expression changes in two Arabidopsis ecotypes and their reciprocal hybrids. Plant Cell 24: 875–892

Shivaprasad PV, Dunn RM, Santos BA, Bassett A, Baulcombe DC (2012) Extraordinary transgressive phenotypes of hybrid tomato are influenced by epigenetics and small silencing RNAs. EMBO J 31: 257–266

Si L, Chen J, Huang X, Gong H, Luo J, Hou Q, Zhou T, Lu T, Zhu J, Shangguan Y, Chen E, Gong C, Zhao Q, Jing Y, Zhao Y, Li Y, Cui L, Fan D, Lu Y, Weng Q, Wang Y, Zhan Q, Liu K, Wei X, An K, An G, Han B (2016) OsSPL13 controls grain size in cultivated rice. Nat Genet 48: 447–456

Springer NM, Stupar RM (2007) Allelic variation and heterosis in maize: how do two halves make more than a whole? Genome Res 17: 264–275

Tan F, Lu Y, Jiang W, Wu T, Zhang RY, Zhao Y, Zhou DX (2018) DDM1 Represses Noncoding RNA Expression and RNA-Directed DNA Methylation in Heterochromatin. Plant Physiology 177: 11871197

Tan F, Zhou C, Zhou Q, Zhou S, Yang W, Zhao Y, Li G, Zhou DX (2016) Analysis of Chromatin Regulators Reveals Specific Features of Rice DNA Methylation Pathways. Plant Physiol 171: 2041–2054

Wendte JM, Zhang Y, Ji L, Shi X, Hazarika RR, Shahryary Y, Johannes F, Schmitz RJ (2019) Epimutations are associated with CHROMOMETHYLASE 3-induced de novo DNA methylation. Elife 8

Xie F, Zhang J (2018) Shanyou 63: an elite mega rice hybrid in China. Rice (N Y) 11: 17

Xie W, Wang G, Yuan M, Yao W, Lyu K, Zhao H, Yang M, Li P, Zhang X, Yuan J, Wang Q, Liu F, Dong H, Zhang L, Li X, Meng X, Zhang W, Xiong L, He Y, Wang S, Yu S, Xu C, Luo J, Li X, Xiao J, Lian X, Zhang Q (2015) Breeding signatures of rice improvement revealed by a genomic variation map from a large germplasm collection. Proc Natl Acad Sci U S A 112: E5411–5419

Yan W, Liu H, Zhou X, Li Q, Zhang J, Lu L, Liu T, Liu H, Zhang C, Zhang Z, Shen G, Yao W, Chen H, Yu S, Xie W, Xing Y (2013) Natural variation in Ghd7.1 plays an important role in grain yield and adaptation in rice. Cell Res 23: 969–971

Yan WH, Wang P, Chen HX, Zhou HJ, Li QP, Wang CR, Ding ZH, Zhang YS, Yu SB, Xing YZ, Zhang QF (2011) A Major QTL, Ghd8, Plays Pleiotropic Roles in Regulating Grain Productivity, Plant Height, and Heading Date in Rice. Molecular Plant 4: 319–330

Yu SB, Li JX, Xu CG, Tan YF, Gao YJ, Li XH, Zhang Q, Saghai Maroof MA (1997) Importance of epistasis as the genetic basis of heterosis in an elite rice hybrid. Proc Natl Acad Sci U S A 94: 9226–9231

Zemach A, Kim MY, Hsieh PH, Coleman-Derr D, Eshed-Williams L, Thao K, Harmer SL, Zilberman D (2013) The Arabidopsis nucleosome remodeler DDM1 allows DNA methyltransferases to access H1-containing heterochromatin. Cell 153: 193–205

Zemach A, Kim MY, Silva P, Rodrigues JA, Dotson B, Brooks MD, Zilberman D (2010) Local DNA hypomethylation activates genes in rice endosperm. Proc Natl Acad Sci U S A 107: 18729–18734

Zhang J, Chen LL, Xing F, Kudrna DA, Yao W, Copetti D, Mu T, Li W, Song JM, Xie W, Lee S, Talag J, Shao L, An Y, Zhang CL, Ouyang Y, Sun S, Jiao WB, Lv F, Du B, Luo M, Maldonado CE, Goicoechea JL, Xiong L, Wu C, Xing Y, Zhou DX, Yu S, Zhao Y, Wang G, Yu Y, Luo Y, Zhou ZW, Hurtado BE, Danowitz A, Wing RA, Zhang Q (2016) Extensive sequence divergence between the reference genomes of two elite indica rice varieties Zhenshan 97 and Minghui 63. Proc Natl Acad Sci U S A 113: E5163–5171

Zhang M, Xie S, Dong X, Zhao X, Zeng B, Chen J, Li H, Yang W, Zhao H, Wang G, Chen Z, Sun S, Hauck A, Jin W, Lai J (2014) Genome-wide high resolution parental-specific DNA and histone methylation maps uncover patterns of imprinting regulation in maize. Genome Res 24: 167–176

Zhang Q, Li Y, Xu T, Srivastava AK, Wang D, Zeng L, Yang L, He L, Zhang H, Zheng Z, Yang DL, Zhao C, Dong J, Gong Z, Liu R, Zhu JK (2016) The chromatin remodeler DDM1 promotes hybrid vigor by regulating salicylic acid metabolism. Cell Discov 2: 16027

Zhang Q, Wang D, Lang Z, He L, Yang L, Zeng L, Li Y, Zhao C, Huang H, Zhang H, Zhang H, Zhu JK (2016) Methylation interactions in Arabidopsis hybrids require RNA-directed DNA methylation and are influenced by genetic variation. Proc Natl Acad Sci U S A 113: E4248–4256

Zhou G, Chen Y, Yao W, Zhang C, Xie W, Hua J, Xing Y, Xiao J, Zhang Q (2012) Genetic composition of yield heterosis in an elite rice hybrid. Proc Natl Acad Sci U S A 109: 15847–15852

Zhou Q, Lim JQ, Sung WK, Li G (2019) An integrated package for bisulfite DNA methylation data analysis with Indel-sensitive mapping. BMC Bioinformatics 20: 47

